# Felidae call type and species identification based on acoustic features

**DOI:** 10.1101/2022.03.30.486147

**Authors:** Danushka Bandara, Karen Exantus, Cristian Navarro-Martinez, Murray Patterson, Ashley Byun

## Abstract

The cat family Felidae is one of the most successful carnivore lineages today. However, the study of the evolution of acoustic communication between felids remains a challenge due to the lack of fossils, the limited availability of audio recordings because of their largely solitary and secretive behavior, and the underdevelopment of computational models and methods needed to address acoustic evolutionary questions. This study is a first attempt at developing a machine learning-based approach to the classification of felid calls as well as the identification of acoustic features that distinguish felid call types and species from one another. A felid call dataset was developed by extracting audio clips from diverse sources. The audio clips were manually annotated for call type and species. Due to the limited availability of samples, this study focused on the Pantherinae subfamily. Time-frequency features were then extracted from the Pantherinae dataset. Finally, several classification algorithms were applied to the resulting data. We achieved 91% accuracy for this Pantherinae call type classification. For the species classification, we obtained 86% accuracy. We also obtained the most predictive features for each of the classifications performed. These features can inform future research into the evolutionary acoustic analysis of the felid group.

## I. INTRODUCTION

Acoustic communication plays a critical role in many tetrapod species (Chen and Weins et al. 2020). Such vocalizations can transmit objective information about an individual’s internal state as well as information critical for intraspecific interactions such as mate selection and territorial defense (Hauser, 1996; Peters and Peters 2010; Tavernier et al. 2020).

The cat family Felidae represents one of the most successful carnivore lineages today with as many as 40 recognized species organized into eight major lineages (Pecon-Slattery et al. 2004). Despite the fact that most adult felids are largely solitary (Kitchener 1991), vocalization still plays a critical role in intraspecific communication (Peters 1978). Current knowledge suggests that all felid species share the same fundamental call repertoire (Peters, 1991). There are 14 major discrete and graded calls documented in Felidae (Peters, 1991; Sunquist and Sunquist, 2002). These calls vary in three major structural domains: loudness (amplitude), time (call duration), and pitch (frequency), though there is also considerable variability in other acoustic features such as harmonic structure. Learning does not seem to play a significant role in felid vocalizations. Rather, the species-specific acoustic structure of these calls is largely genetically determined (Peters, 1978; Peters and Tonkin-Leyhausen, 1999).

While attempts to understand the evolution of vocalizations have been made within Felidae (see Peters et al. 1999; Peters and Peters 2010), such studies are challenging to conduct due to 1) the lack of fossils that would provide anatomical information about sound production and the acoustic characteristics of ancestral vocalizations, 2) the paucity and the heterogeneity of audio recordings of vocalizations for Felidae species due in part to their largely solitary and secretive behavior, and 3) the underdevelopment of computational models and methods needed to address acoustic evolutionary questions.

This study uses validated machine learning methods to address the aforementioned second and third difficulties. We focused specifically on vocalizations from the subfamily Pantherinae which includes species such as lions, tigers and jaguars, due to the larger number of audio files that could be obtained. This study curates calls from various sources into a single uniform annotated database. This allows standardization for preprocessing and model building. The heterogeneity of audio recordings of the calls is addressed by converting the raw audio into time domain, time-frequency domain, cepstral, and statistical features. Since these features describe the salient characteristics of the auditory signals, they enable combining raw audio files from diverse sources. The development of machine learning models provides an automated methodology to classify such audio calls without human judgment. Machine learning models also provide insight into the important audio features that characterize the different vocalizations in different species. Such identification and extraction of such features will help promote further research of animal vocalizations and the evolution of acoustic communication.

### A. Feature extraction methods used in felid call classification

Previous research into animal call identification/classification has broadly considered several groups of acoustic features:

1. Time-frequency domain features (Nanni et al., 2020; Nanni et al., 2020; Raccagni & Ntalampiras, 2020)
2. Cepstral features (Balemarthy et al., 2018; Kukushkin & Ntalampiras, 2021; Ntalampiras et al., 2019). Cepstral features are a well-understood methodology (Davis & Mermelstein, 1980.) for representing audio signals in terms of their short time power spectra.
3. Deep learning methods: Several newer studies use deep neural networks to achieve felid sound classification (e.g. Pandeya & Lee., 2018; Pandeya et al., 2018). However, the drawback of these methods is their ‘black box’ nature. These methods, by default, do not provide interpretable acoustic features that can be used for further analysis, such as evolutionary acoustic analysis. Also, these methods require many samples, which is challenging regarding certain felid species.

### B. Models used in felid call classification

Classification of felid calls has also been attempted in several studies, mainly as part of a larger dataset in combination with other animal species calls.

For example, Weninger, F., & Schuller, B. (2011) developed a classification method by left-right and cyclic Hidden Markov Models, recurrent neural networks with Long Short-Term Memory, and Support Vector Machines, achieving up to 81.3 % accuracy on a 2-class, and 64.0 % on a 5-class classification task. Ji et al. (2013) achieved individual animal identification using a hidden Markov model (HMM) with frame-based spectral features consisting of cepstral coefficients.

In our study, we developed a five class call classification and four class species classification. The classifiers used frequency based features as well as time based features without relying on state based features. Leaving out state based features in favor of simpler features allowed simpler preprocessing and less likelihood of overfitting.

### C. Goals of this study

In summary, there are three main goals of this study,

1. To develop an annotated felid call dataset by combining audio from diverse sources.
2. To objectively detect key acoustic features that distinguish between types of vocalizations and identify species-specific differences.
3. To develop machine learning classifiers that automate felid call identification.

## II. METHODS

### A. Data collection

One major challenge this study faces is the lack of a standardized database. The call data used in this study were collected by different researchers for different intended applications. Some other calls were extracted from online videos that were ambient recordings that contain a felid vocalization as part of a longer recording. The dataset that we use in this study was collected from various sources, including Animal Sound Archive (Museum für Naturkunde Berlin), calls provided by G. Peters, and social media (see Table I).

**TABLE I.** Sources of felid call audio clips.

|  | Leopard | Clouded Leopard | Lion | Jaguar | Snow Leopard |
| --- | --- | --- | --- | --- | --- |
| Museum für Naturkunde | 0 | 0 | 0 | 4 | 0 |
| Youtube | 14 | 0 | 9 | 20 | 10 |
| TikTok | 0 | 12 | 0 | 0 | 0 |
| G. Peters | 1 | 0 | 1 | 1 | 0 |
| Facebook | 0 | 2 | 0 | 0 | 2 |

These latter sound files were downloaded and converted to WAV file format from MP4. Following the conversion, all sound files were manually annotated using Raven Pro, GarageBand, and Melodyne 5, which all assisted in identifying the specific calls. These manual annotations were saved into individual clips of the complete sound files labeled as the type of felid call. A metadata file was connected with each audio sample, including file name, the onset of the call, duration of the call, species name, gender, age, audio quality, and any relevant notes.

The distribution of the raw audio files by Pantherinae lineage and vocalizations are provided in Table II.

**TABLE II.** Contents of the felid call dataset.

|  | Leopard | Clouded Leopard | Lion | Jaguar | Snow Leopard |
| --- | --- | --- | --- | --- | --- |
| hiss | 0 | 6 | 0 | 0 | 0 |
| prusten | 0 | 29 | 0 | 11 | 4 |
| roar | 29 | 0 | 25 | 11 | 0 |
| growl | 12 | 0 | 8 | 0 | 0 |
| puff | 4 | 0 | 1 | 0 | 0 |
| mew | 0 | 0 | 0 | 0 | 3 |

**TABLE III:** Classification accuracies for call and species identification using various classification algorithms.

| Model | SVM | XGBoost | LightGBM | Logistic Regression | Random Forest |
| --- | --- | --- | --- | --- | --- |
| Call Classification Accuracy | 81% | 86% | 86% | 91% | 88% |
| Species Classification Accuracy | 84% | 86% | 81% | 79% | 84% |

Since certain species of cats are both rare and elusive, obtaining a substantial number of audio samples for those species remains a challenge. Thus, the obtained dataset has a class imbalance. The snow leopard species and mew calls were dropped from the analysis because there weren’t enough samples to include in training and test sets to provide meaningful results.

### B. Signal preprocessing

After the dataset development, the next step was signal preprocessing. This step involved the conversion of the waveform from the time domain to the frequency(spectrum) and to the time-frequency domain(spectrogram).

### C. Acoustic feature extraction

Below is a list of features extracted and used for the machine learning models. We used the mean of several of these features in our model, as is common in audio processing. Studies have shown that taking the mean does not always result in the loss of necessary information (Rana & Jain, 2014).

Frequency Range - A continuous range or spectrum of frequencies that extend from one limiting frequency to another.

Amplitude Range - The difference between the highest positive and the lowest positive amplitude.

Average Amplitude - The mean of the amplitudes.

Root-mean-square (RMS) energy – A measure of the audio file’s loudness in energy per frame.

Pulses per Sec - This feature is calculated as the number of peaks per second of the audio clip.

Duration - The difference between start time and end time from the audio file.

Zero-Crossing - The number of signal changes between negative and positive values in the spectrum.

Mel-spectrogram - Mel frequency refers to a mapping of audio from the linear frequency domain to the Mel frequency domain. The Mel frequency domain mimics human perception of audio. The Mel spectrogram is the spectrogram of the audio over the Mel scale. This feature consists of the average value of the Mel spectrogram.

Mel frequency cepstral coefficients(MFCC)-MFCCs are obtained by taking the Mel spectrogram and applying a discrete cosine transformation. The 20 such MFCCs are averaged to obtain this feature.

Spectral Rolloff - This feature computes the rolloff frequency (The spectrogram bin such that at least 85% of the energy of the spectrum in this frame is contained in this bin and the bins below) for each frame in a signal. We used the mean value of the array of frequencies.

Spectral Contrast - Spectral contrast considers the spectral peak, the spectral valley, and their difference in each frequency subband. We used the mean of these values.

Chromagram - Chromagram is a variation on time-frequency distributions, representing spectral energy. It is computed from a waveform. We used the mean value.

Tempogram-It is a local autocorrelation of the onset strength envelope. The tempogram computes tempo variation and local pulse in the audio signal. We used the mean value.

### D. Data cleaning

The data was then analyzed to check for any null values or outliers that may have resulted from the extraction stage and cleaned to prepare for further analysis of the feature values. After the data was cleaned, the features made up of integers and floating-point values were standardized to better fit the models.

### E. Classification

Once the data was cleaned and ready to be used in the models, the features and the labels (call or species) progressed to the classification stage. The data was then split 70/30, with 70% of the data being used to train the model and 30% being used to test the model.

The classification models used include:

Support vector machines (SVM): The SVM algorithm attempts to find an optimal hyperplane that separates the training data in euclidean space. This is achieved by maximizing the margin between two hyperplanes that lie on the support vectors(the data points that belong to two separate classes and are ‘closest’ to each other)

Random forest: Random forests are formed by multiple decision trees. Decision trees in turn are models that split the dataset at multiple levels, choosing the split with the highest ‘information gain’ at each level.

Gradient boosting: Gradient boosting is a modification of the random forest algorithm such that the trees each learn sequentially. I.e. each tree learns the ‘residual’ from the previous step. XGBoost and LightGBM both use this method with different implementations.

Logistic regression: Logistic regression model learns a logistic function that separates the data points into classes. The model is optimized using gradient descent applied to the loss function (difference between output of model and the actual label).

## III. RESULTS

The goal of this research was to develop models for felid call and felid species classification. In section II part a, we described how the dataset of felid calls was created using diverse sources. In this section, we present the results of the machine learning classification.

### B. Call classification

The best classification accuracy for call classification was from the logistic regression classifier. The confusion matrix and most predictive features for this classifier is given in figure 3.

**FIG. 1.**
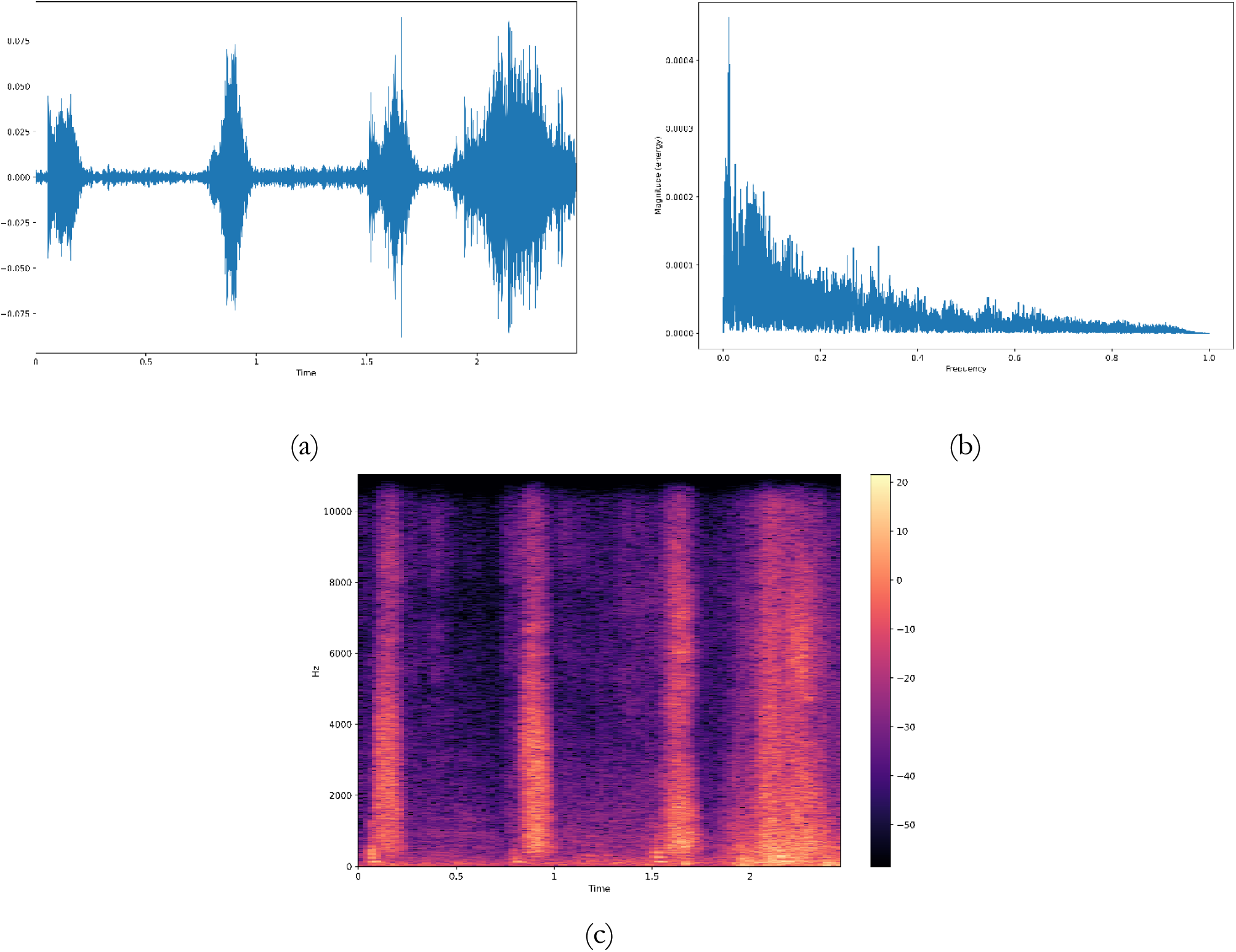
(a) Puff waveform from a lion (b) Puff spectrum from a lion (c) Puff spectrogram from a lion

**FIG. 2.**
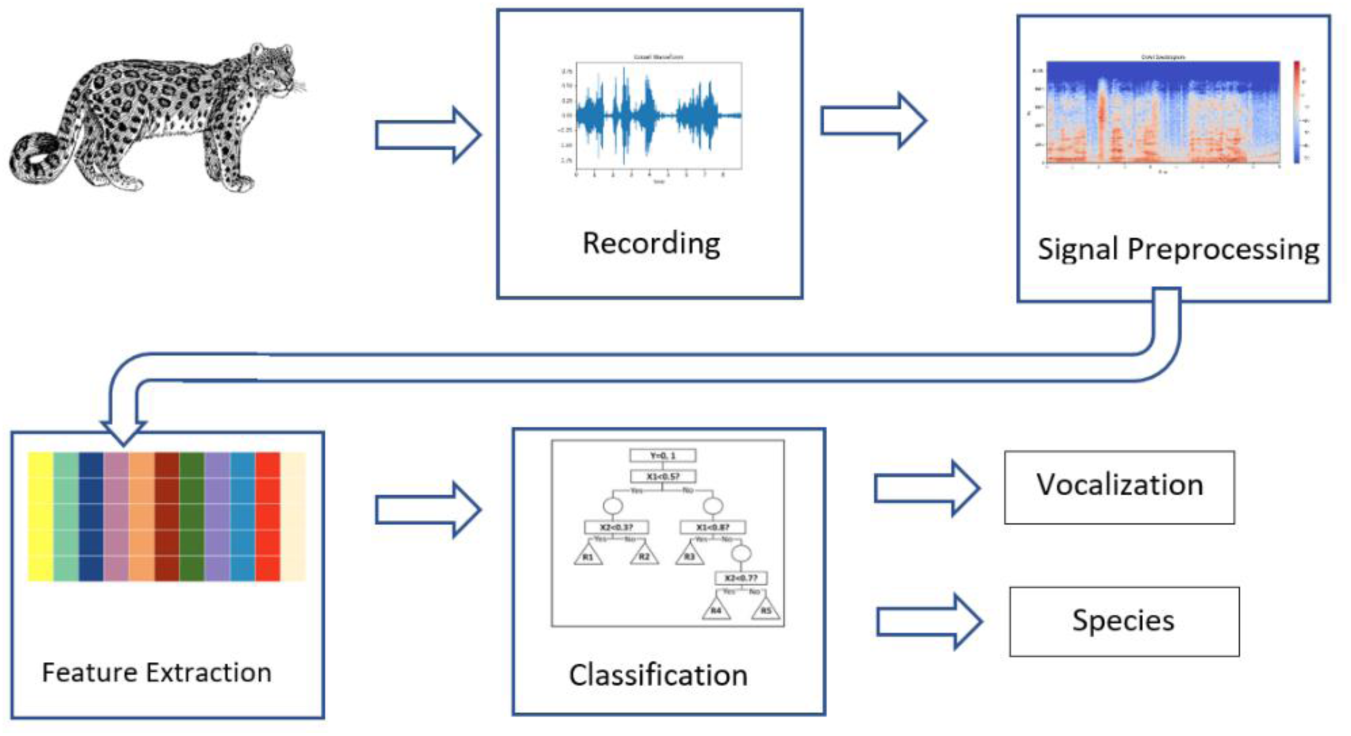
Flowchart of the machine learning classification of species and vocalizations.

**FIG. 3.**
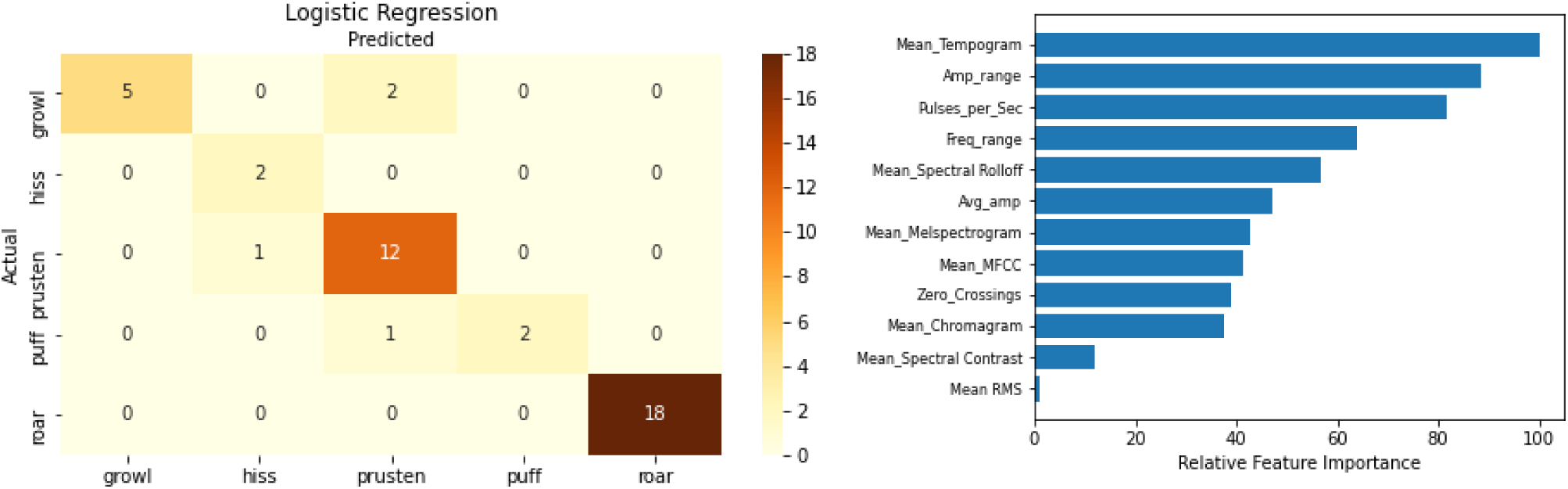
Confusion matrix and feature importance for the logistic regression call classifier.

The receiver operating characteristic (ROC) curve shows the tradeoff between true positive rate and false positive rate. Since the ROC curve does not depend on the class distribution, it is useful for evaluating performance in classifications where there is a class imbalance. The ROC curve for the call classification is given in figure 4.

**FIG. 4.**
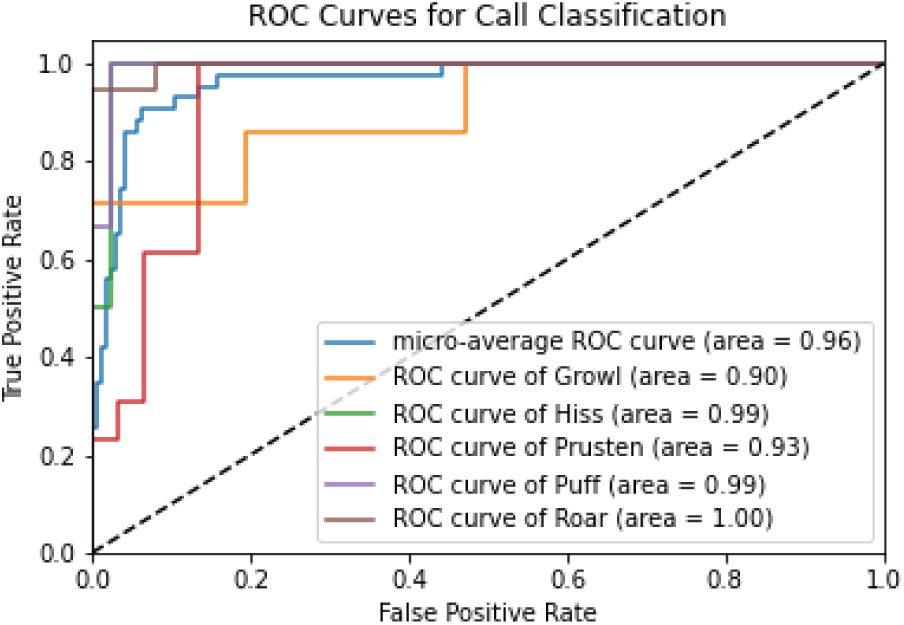
ROC curves for the logistic regression call classifier.

The area under the ROC curve (AUC) is a measure of classifier performance. From figure. 4 we can see that the area under the curve values for all the call types are at or above 0.9.

### B. Species classification

The best classification performance for species classification was obtained by the XGBoost classifier. The confusion matrix and feature importance for the classifier are given in figure 5.

**FIG. 5.**
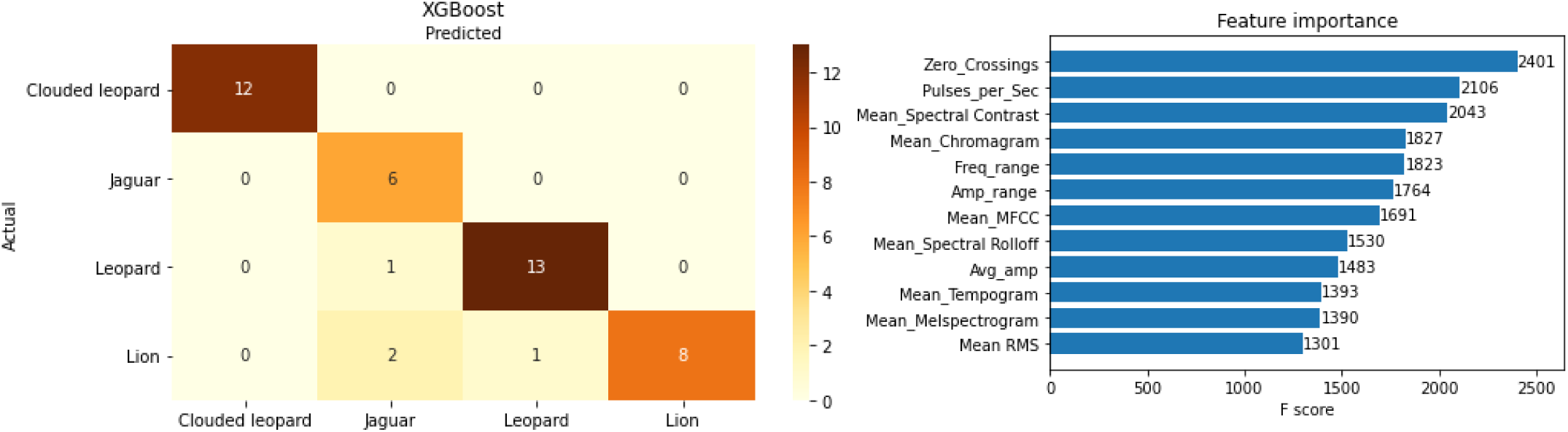
Confusion matrix and feature importance for the XGBoost species classifier.

From figure 6 we can see that the area under the ROC curve for the species classifier is at or above 0.84 for all species. The worst AUC of 0.84 was for the lion species.

**FIG. 6.**
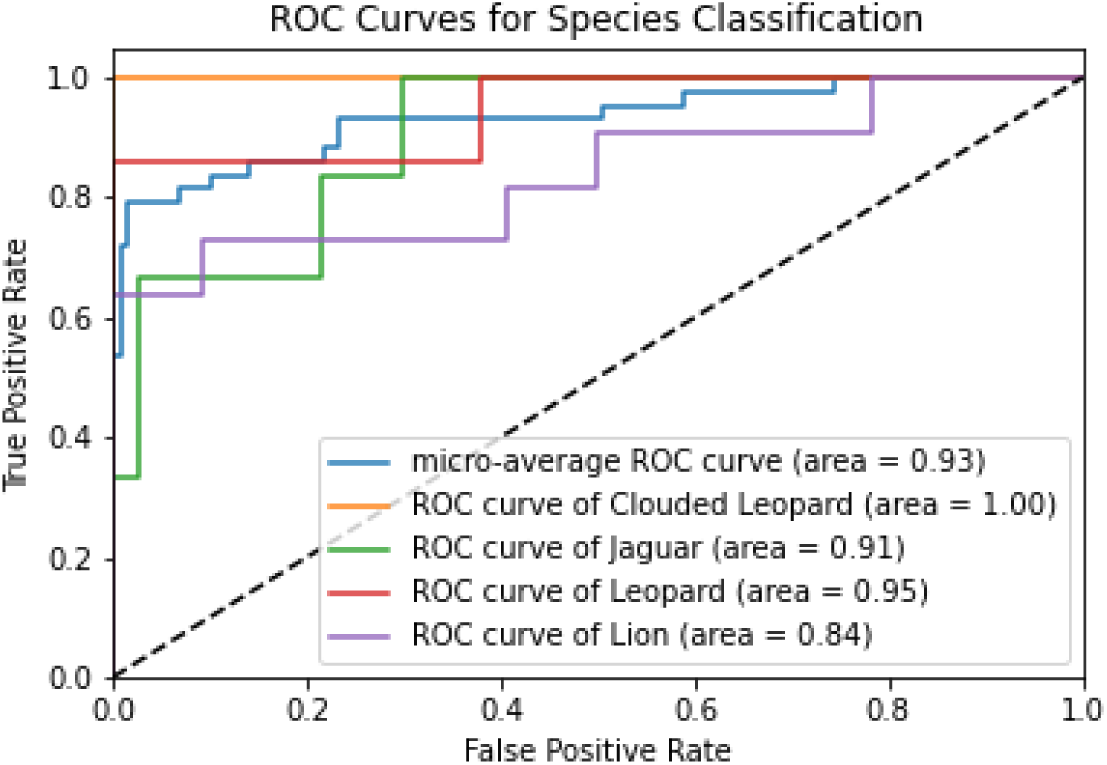
ROC curves for the XGBoost species classifier.

## IV. DISCUSSION

We found that zero crossings, mean tempogram, frequency range, amplitude range and mean MFCC features appear to be successful in both call and species classifications For the call classifier, growl and prusten have the lowest area under the curve. This is evident from looking at the confusion matrix where growl and prusten accounted for 3 of the 5 misclassifications. For the species classification, lion species had the lowest area under the curve. Similarly, lion species accounted for 3 of the 4 misclassifications.

Overall, our classifiers were able to obtain better classification performance than comparable felid call classification studies. (Weininger et al., 2011).

## V. CONCLUSION

This research aimed to develop a standardized dataset of felid calls and build an automated model for felid calls and species classification. The most predictive features were identified and reported for use in future research. The paper also explained the preprocessing, feature engineering, data cleaning, and machine learning steps. Several models were fitted to the data, and the best performing were Logistic Regression for call classification and XGBoost for species classification, obtaining 91% and 86% accuracy, respectively. These results show that we can classify felid calls by vocalization type or species using only an audio recording of their call. Our results support the claim that felid vocalizations share certain features that are universally applicable and genetically based as opposed to learned behavior.

This is an emerging area of research, and there is a lack of readily available datasets. Therefore, having a sizable standardized felid call dataset will help future researchers develop better classifiers. The dataset we have built is an initial attempt at this. We did not consider where the audio was collected and other relevant metadata. Incorporating those and collecting more call data could help get better classification performance.

## ACKNOWLEDGMENTS

The authors would like to acknowledge the contributions by: Sai Greeshma Saladi, Rohindraj Kandasamy, Nicholas Furey, Tianyu Yang to the data preprocessing and Brendan Smith, Rebecca Buopane for assisting in audio annotations. We are grateful for the audio files provided by Dr. Gustav Peters, former Curator of Mammals, Zoological Research Museum, Bonn and Dr. Karl-Heinz Frommolt, Scientific Head of the Animal Sound Archive (Museum für Naturkunde Berlin). This work was supported by the Fairfield University Science Institute (AB and MP), the Fredrickson Family Innovation Lab Grant (AB and MP) and Georgia State University Computer Science Start Up Grant (MP).

## Notes

### Competing Interest Statement

The authors have declared no competing interest.

